# Open neuroinformatics infrastructure ecosystem for federated multisite studies

**DOI:** 10.64898/2026.04.30.721944

**Authors:** Michelle Wang, Nikhil Bhagwat, Francesco Cremonesi, Mathieu Dugré, Julia-Katharina Pfarr, Emile d’Angremont, Alyssa Dai, Arman Jahanpour, Sebastian Urchs, Sergen Cansiz, Lucie Chambon, Ali Tolga Dinçer, Jhonatan Torres, Marc Vesin, Gabriel Pinilla-Monsalve, Yuan Song, Chris Vriend, Francis Jeanson, Oury Monchi, Ysbrand D. van der Werf, Marco Lorenzi, Jean-Baptiste Poline

## Abstract

Despite growing understanding of the benefits of having Findable, Accessible, Interoperable, and Reusable (FAIR) data, many datasets still cannot be shared. Federated analysis methods can enable multisite studies that do not require the sharing of participant-level information. However, there are many practical hurdles that prevent the large-scale adoption of federated methods. We discuss challenges related to cross-site data preparation for federated learning, present solutions offered by recent neuroinformatics projects, and showcase an example of tool integration applied to neurodegenerative disease data.

## Main text

The field of neuroimaging is seeing an ever-growing number of datasets. However, constraints^1^ driven by e.g., increasingly stringent data privacy regulations^2,3^ have led to many datasets not being shared^4,5^: they remain in so-called “silos”, which prevents reuse, contributes to the ongoing reproducibility crisis^6^, and impedes scientific discoveries^7^.

Because current global trends are moving towards an increase rather than a decrease of data-sharing constraints, there have been recent efforts in developing methods that enable multisite studies while respecting data-sharing limitations. Notably, federated learning^8,9^ is a machine learning paradigm where models are fitted on data from different sites without the need to share those data. Recently, several software frameworks for federated learning have emerged, including NeuroFLAME (formerly COINSTAC)^10^ and Fed-BioMed^11^ in healthcare, as well as domain-agnostic frameworks such as Flower^12^ and NVIDIA FLARE^13^. However, despite the maturity of these projects, there has been limited real-world application of federated learning in neuroimaging so far: a non-systematic PubMed search conducted in March 2026 using the keywords [“Federated learning” or “Federated analysis”] and “Neuroimaging” found only 4 papers^14–17^ that were not review articles or method development articles using simulated federated setups.

The unfortunate reality is that the technical barrier of entry to federated analyses is still too high for the average neuroimaging researcher. A multi-site federated experiment requires each site to 1) set up the necessary local infrastructure required by a given federated learning framework and 2) prepare data in a way that is identical to every other site, so that all datasets can be readily used by the framework. Both of these requirements can present technical hurdles, but the latter, data preparation, is particularly challenging and often overlooked. In practice, it is difficult to ensure that neuroimaging datasets at different sites have been processed and formatted in exactly the same way, especially when under data-sharing constraints. We highlight three main challenges here: filesystem organization, data processing, and variable semantics (Fig. 1a). All of these are essential for federated analyses and typically require significant manual work that is difficult to coordinate between sites. Although efforts exist to address individual components, there remains a lack of integrative solutions that connect all aspects of data preparation into a streamlined, easy-to-follow workflow.

**Figure 1:**
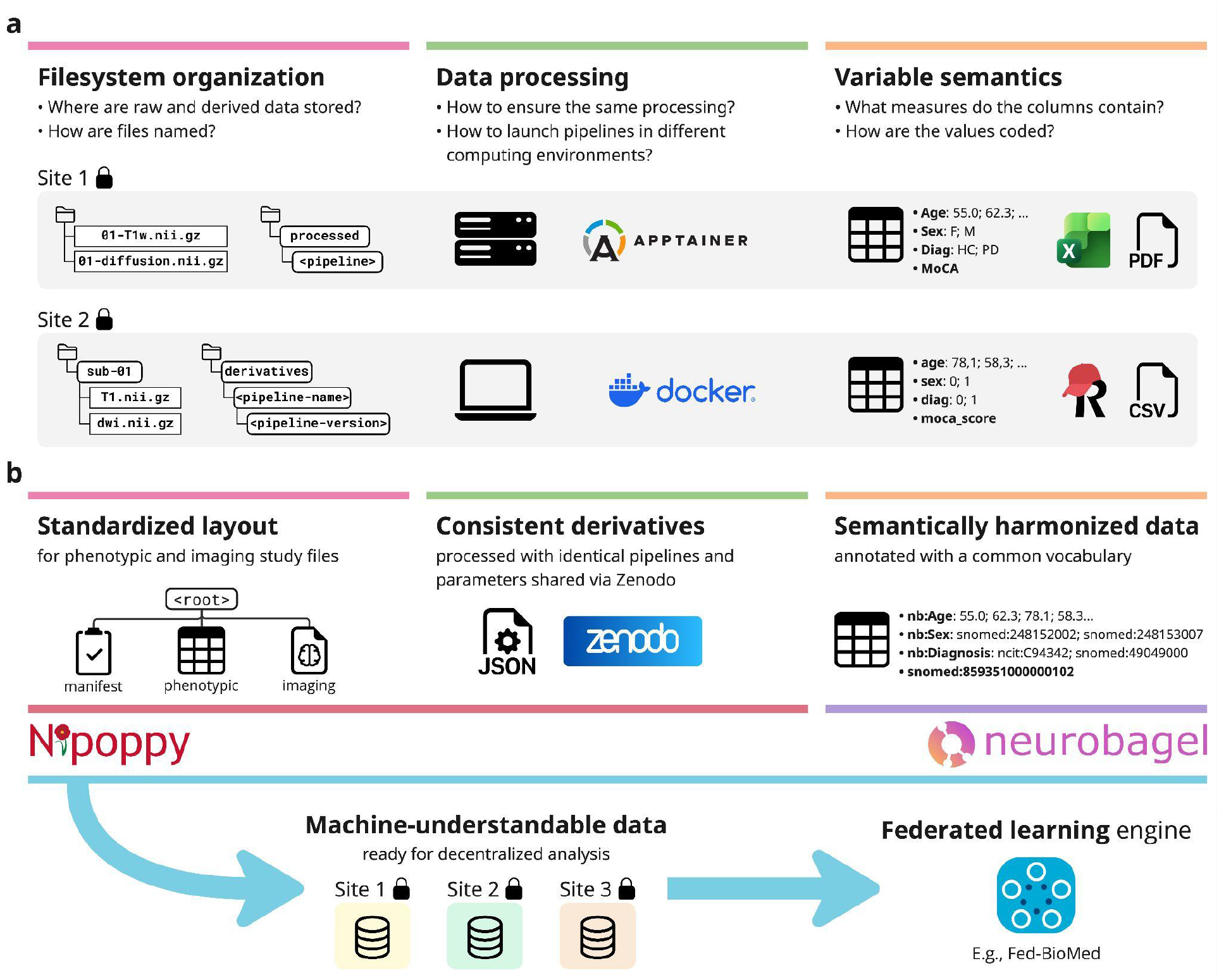
Challenges and solutions related to data preparation for federated learning analyses. **a**: We identify filesystem organization, data processing, and variable semantics as key but often overlooked challenges that increase the barrier of entry to federated learning. **b**: These challenges can be addressed through the standardization of filesystem layout, consistent and replicable processing of data, and semantic harmonization of variables. The Nipoppy and Neurobagel projects provide tools that implement these solutions and facilitate the creation of machine-understandable, analysis-ready data to be used as input in federated analyses.

We propose an integrative ecosystem that addresses these three challenges, specifically using the Nipoppy framework for standardized organization and processing of neuroimaging-clinical datasets^18^ and the Neurobagel ecosystem for distributed dataset harmonization and cohort discovery^19^ (Fig. 1b). We show how tools from these projects can help create standardized machine-understandable, analysis-ready data to be used in federated analyses. We further describe the integration of these tools with the Fed-BioMed framework^11^ to enable seamless definition and launching of new federated learning experiments without any additional setup at local sites. Finally, we showcase a practical example where this integration is leveraged to conduct an analysis comparing federated and centralized setups in predicting age, cognitive decline and diagnosis.

### Challenge 1: Filesystem organization

Different research groups rarely follow the same directory and file structure to organize study data, which makes it difficult for both humans and machines to know where different types of data are located. The clear solution to this challenge is to adopt a consistent data organization standard, such as the Brain Imaging Data Structure (BIDS)^20^, across sites. We argue for the importance of a data organization standard at the *entire study level*, as analyses often require multiple types of data (e.g. raw and derived, imaging and non-imaging). The BIDS standard has recently been extended to include a study-level specification, which aims to encompass more than just raw or derived imaging data. A stricter (but fully compliant) version of this specification is prescribed by the Nipoppy framework. It encompasses all of a study’s data and describes where and how raw imaging data, derived imaging data and tabular phenotypic data should be stored, allowing for the automated generation of machine-understandable, analysis-ready data.

### Challenge 2: Data processing

Most neuroimaging analyses require imaging data to be at least minimally preprocessed (e.g., brain extraction, intensity normalization). Often, machine learning or statistical models are applied not on the images themselves, but rather on features extracted from them (sometimes called “imaging-derived phenotypes”, or IDPs). The choice of processing pipeline and version can greatly affect the distribution of IDPs^21,22^, and so it is critical to ensure that all sites participating in a federated analysis use the same pipelines, including version and runtime parameters. Portability of software environments can be achieved through the use of containerization technology^23–25^, and programmatic launching or re-launching of pipelines can be achieved with tools and frameworks such as the DataLad distributed data management system^26^ and the Boutiques command-line description framework^27^. The Nipoppy framework leverages Boutiques to enable consistent and decentralized processing of neuroimaging data. Pipeline configuration files specifying software version and runtime parameters are shared via the Nipoppy pipeline store^28^ (on the Zenodo data repository) and can be downloaded locally for each dataset with a user-friendly command-line interface. This enables researchers coordinating a federated analysis to easily share the exact container image and runtime parameters to be used, ensuring identical data processing across sites.

### Challenge 3: Variable semantics

Different studies have their own ways of collecting phenotypic data (demographics, assessment scores, etc.), often with unique terminologies and encoding schemes for variable names and values. Semantic harmonization of these vocabularies (i.e., alignment to a standardized vocabulary such as an established ontology) is needed to enable data from different sites to be analyzed together. The creation of a machine-understandable file that maps study-specific names to standardized terms is facilitated by the user-friendly Neurobagel annotation tool (https://annotate.neurobagel.org/). When such a mapping has been done by all sites participating in a federated analysis, harmonized views of these datasets can be created and combined in a joint analysis.

Study data that are organized according to the Nipoppy specification and annotated with Neurobagel can easily be understood at the filesystem (data organization) and semantic (data content) levels, and can be processed with identical pipelines and parameters using Nipoppy tools. We leveraged this standardization to facilitate federated learning experiments that use the Fed-BioMed framework. Using data from six neuroimaging studies involving participants with Parkinson’s (PD) or Alzheimer’s disease (AD), all prepared in this way, we compared a Fed-BioMed federated analysis setup with two traditional centralized experimental setups – siloed (no sharing) and mega-(central pooling of data) analyses (Fig. 2a). We ran three experiments based on different machine learning tasks: predicting age, future cognitive decline (within 5 years from baseline), and diagnosis (PD/AD patient vs control) from structural brain measures and demographic variables. We find that models trained with the Siloed setup do not generalize well to other datasets (Fig. 2b), and that the average model performance is better for Federated compared to Siloed and similar between the Federated and Mega-analysis setups (Fig. 2c). The dataset standardization achieved through Nipoppy and Neurobagel greatly streamlined this analysis, since new experiments could be launched without requiring any changes to local datasets (see Methods); in contrast, data preparation without such standardization would have required many custom, dataset-specific operations, increasing the workload significantly.

**Figure 2:**
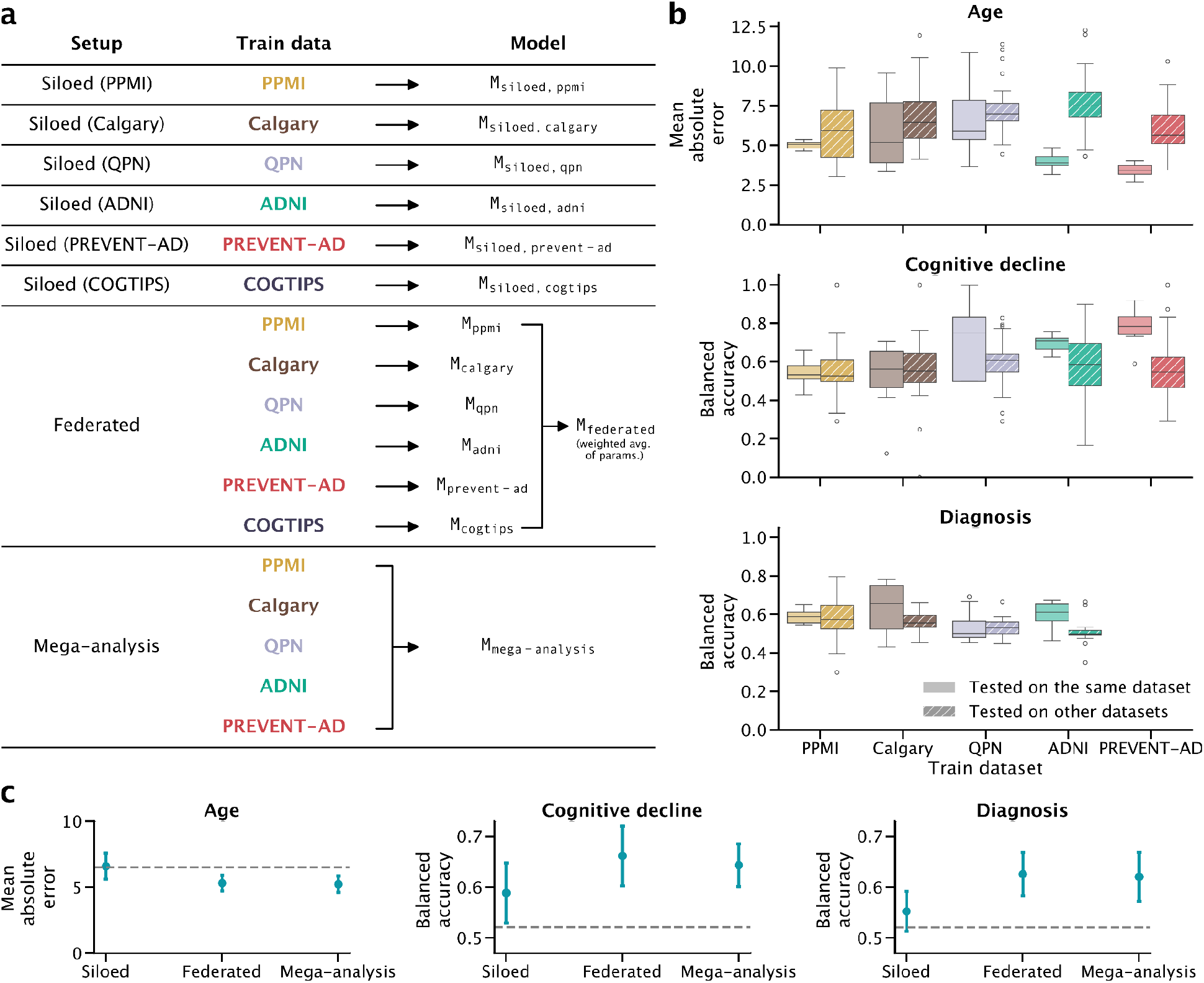
Experiment with neuroimaging prediction tasks under different data-sharing constraints. **a**: Machine learning models and data-sharing setups. **b**. Performance of Siloed models tested on in- or out-of-sample data. **c**: Model performance comparison between setups. Error bars represent 1 standard deviation. Dashed lines represent the 5th, 95th and 95th percentiles of the null model distributions for the Age, Cognitive decline and Diagnosis tasks respectively.

The growing ecosystem of neuroinformatics infrastructure for decentralized data processing and IDP extraction (supported by the Nipoppy framework) and harmonization (enabled by the Neurobagel project) can facilitate the implementation of federated analyses, allowing researchers to focus on scientific questions instead of technical setup. Furthermore, the use of previously unavailable data can foster more inclusive research and drive the development of reliable and generalizable models. Our experiments on age, diagnosis and cognitive decline prediction in different machine learning setups show that models trained in a federated setup perform better than models trained on siloed datasets, further cementing the notion that federated learning is a worthwhile analysis option when datasets cannot be directly shared. We hope these results, combined with our work on decentralized neuroimaging infrastructure, can precipitate a paradigm shift in neuroimaging research that leads to future large, multisite neuroimaging studies that maintain full local data governance.

## Online methods

Below are detailed methods for the analysis in which we leveraged tools from the Nipoppy and Neurobagel projects to conduct a federated analysis in which we compared Siloed, Federated and Mega-analysis setups in machine learning tasks applied to neurodegenerative disease data. Custom code developed specifically for this analysis is openly available at https://github.com/neurodatascience/fl-pd/.

### A federated learning framework with Nipoppy, Neurobagel and Fed-BioMed

A typical federated learning setup involves a network of sites that communicate via client-server protocols to launch experiments and update model weights. To train a model, a designated server node defines and disseminates an experiment plan to client nodes, where datasets are hosted; this plan details input and output features and model choices. Model fitting occurs locally at each client node, and a global model is updated iteratively on the server side. The researcher coordinating the federated experiment manages the server and sets experiment details. The data manager at each site is responsible for data preparation: they must ensure that the data added to client nodes meets the expected requirements (processing pipelines, phenotypic column names and/or order, etc.). These requirements are usually communicated separately by the researcher, for example via email. Without adequate standardization methods, local data preparation can be time-consuming and error-prone, as every site will likely need to develop custom solutions for their own dataset. Standardization with Nipoppy and Neurobagel tools can greatly simplify data preparation and reduce the need for dataset-specific wrangling.

First, each site can independently adopt the Nipoppy framework, using standard pipelines to convert raw data to the BIDS standard^20^. Then, the researcher leading the federated experiment can easily share relevant pipeline configuration files to data managers (via the Zenodo platform), allowing them to process raw data and extract IDPs in a decentralized manner with Nipoppy. For phenotypic data, data managers can use the Neurobagel annotation tool to create data dictionaries where study-specific terms for column names and values are mapped to a standardized vocabulary, allowing these data to be harmonized at the semantic level. Thus, Nipoppy and Neurobagel can be used by researchers to produce standardized, machine-understandable data ready for federated analysis.

We now briefly describe the Fed-BioMed framework. In Fed-BioMed, model fitting involves multiple training rounds. During a training round, global model parameters are sent by the server to the clients, which perform training on local data. Then, locally updated model parameters are sent back to the server to be aggregated, most commonly through weighted averaging based on dataset sample sizes^8^. The process is repeated until a stopping criterion is met. Throughout the entire experiment, each site’s dataset stays local and is never shared: only fitted model parameters are shared. In addition, Fed-BioMed uses state-of-the-art security features such as secure aggregation^29^, differential privacy^30^, and homomorphic encryption^31^ to ensure the privacy of local datasets.

In Fed-BioMed, as different experiments and models can require different input data formats, running additional federated learning experiments may require sites to create new files or otherwise modify their datasets. The use of Nipoppy datasets with Neurobagel annotations increases the flexibility of Fed-BioMed experiment definition. Using the `NipoppyDataRetriever` interface added in Nipoppy 0.4.3 combined with the `CustomDataset` interface added in Fed-BioMed 6.2.0, Fed-BioMed clients can retrieve and merge together tabular data files from Nipoppy study directories into harmonized datasets ready for federated learning. For imaging derivatives data, any file that follows the Nipoppy specification for imaging-derived phenotypes (IDPs) can be loaded by the API (https://nipoppy.readthedocs.io/en/latest/explanations/idp_spec.html). For demographic and assessment data, any measure that has Neurobagel annotations can be extracted and transformed (i.e., remapped to a common terminology) accordingly. All tabular data are then merged into a harmonized dataframe that follows the same format for all Nipoppy datasets.

When defining Fed-BioMed experiments, researchers can easily specify the Nipoppy imaging derivatives files (which have standardized paths) and demographic and assessment variables (through standardized vocabulary terms) to use. Defining and launching additional experiments does not require any modifications to the Nipoppy datasets. We are actively contributing this version of the `CustomDataset` back to Fed-BioMed as a `NipoppyDataset` interface.

### Datasets

We used data from six neuroimaging studies of Parkinson’s (PD) or Alzheimer’s disease (AD): the Parkinson’s Progressive Markers Initiative (PPMI, N=1960)^32^, the Alzheimer’s Disease Neuroimaging Initiative (ADNI, N=1106)^33^, the Pre-symptomatic Evaluation of Experimental or Novel Treatments for Alzheimer’s Disease (PREVENT-AD, N=342)^34^, the Quebec Parkinson Network (QPN, N=290)^35^, the Parkinson’s disease and mild cognitive impairment dataset from the University of Calgary (Calgary, N=158)^36^, and the Cognitive Training In Parkinson Study (COGTIPS, N=113)^37^. Basic demographic information is shown in Supplementary Table 1. Data from PPMI, ADNI, PREVENT-AD and QPN were managed in-house, and data from COGTIPS were handled at a different site in Amsterdam. All datasets were organized and processed following the Nipoppy framework, and phenotypic measures were annotated and harmonized using the Neurobagel annotation tool and command-line tool respectively.

### Imaging measures

We processed cross-sectional T1-weighted magnetic resonance images with FreeSurfer^38^ version 7.3.2, which was run as part of fMRIPrep^39^ version 23.1.3^40^ (for PPMI, QPN, and COGTIPS) or 24.1.1^41^ (for ADNI, PREVENT-AD, and Calgary). We used a custom pipeline^42^ to produce tabular analysis-ready files extracted from the FreeSurfer output files.

### Demographic and cognitive measures

Phenotypic measures included biological sex, age, future cognitive decline status, and diagnosis. We defined cognitive decline status using longitudinal information from the Montreal Cognitive Assessment (MoCA) for PD datasets and the Mini-Mental State Examination (MMSE) for ADNI (AD): participants were deemed to exhibit cognitive decline if they showed a rate of change in total MoCA or MMSE score greater than 1 point of decline per year within five years of the imaging visit. For the PREVENT-AD dataset, for which neither assessment was available, participants were classified as exhibiting cognitive decline if they transitioned from “cognitively unaffected” to “mild cognitive impairment (MCI)” without reverting back according to study data. MCI classification was determined in multidisciplinary consensus meetings, based on review of cognitive test result history, blinded to AD biomarkers and genotype.

### Prediction tasks

We chose the following three prediction tasks:

- Prediction of age in healthy controls from biological sex and FreeSurfer subcortical volumes from 17 regions from the default “aseg” atlas^43^. The model used for this task was a scikit-learn SGD regression model with hinge loss via Fed-BioMed’s `FedSGDRegressor` training plan. We used mean absolute error as the evaluation metric for this task.
- Classification of binary cognitive decline status in patients (based on longitudinal MoCA or MMSE scores) from age, biological sex, and mean FreeSurfer cortical thickness measures from 62 Desikan-Killiany-Tourville^44^ regions. The model used for this task was a scikit-learn^45,46^ stochastic gradient descent (SGD) classification model with hinge loss via Fed-BioMed’s `FedSGDClassifier` training plan. We used balanced accuracy as the model evaluation metric for this task. The COGTIPS dataset was not included in this analysis because the time interval between MoCA visits (around 8 weeks) was too short to reliably compute future cognitive decline status.
- Binary classification of diagnosis (AD/PD patient vs. control) from age, biological sex, mean FreeSurfer cortical thickness measures from 62 Desikan-Killiany-Tourville^44^ regions, and FreeSurfer subcortical volumes from 17 regions from the default “aseg” atlas^43^. The model used for this task was a scikit-learn^45,46^ stochastic gradient descent (SGD) classification model with hinge loss via Fed-BioMed’s training plan. `FedSGDClassifier` training plan. We used balanced accuracy as the model evaluation metric for this task.The PREVENT-AD dataset was not included in this analysis due to lack of participant diagnosis information.

Supplementary Figure 1 shows target variable distributions for all three tasks.

### Machine learning setups

We investigated the following three machine learning setups (Fig. 2a), all implemented using Fed-BioMed:

- **Siloed**: The model is trained on a single dataset, with no sharing of data or model parameters. This was achieved in Fed-BioMed by selecting only one of the individual dataset nodes.
- **Federated**: The model is trained via federated learning, where each client node trains a local model, then shares local model parameters to be aggregated into a global model by the researcher server. This was achieved in Fed-BioMed by selecting all of the individual dataset nodes.
- **Mega-analysis**: The model is trained on all of the five locally-available datasets (PPMI, ADNI, PREVENT-AD, QPN, and Calgary) pooled together. This was achieved in Fed-BioMed by having an additional “mega-analysis” node which held the merged dataset.

Model training was done in a completely federated manner, with each dataset node running in an isolated Docker environment (despite some of them running on the same machine). Model evaluation was performed post-hoc using the five datasets that were available locally (PPMI, ADNI, PREVENT-AD, QPN, and Calgary). In all setups, we used 10-fold cross-validation: models were trained on 90% of the data and tested with six test sets: the held-out 10% of each of the five local datasets, as well as on the combination of those held-out subsets. This was done a total of 10 times, with non-overlapping test folds, to obtain an estimate of model generalizability.

### Fed-BioMed deployment

Following instructions on the Fed-BioMed documentation website^47^, we deployed Docker^25^ containers for the server and client processes. A Virtual private network (VPN) server and the Fed-BioMed researcher service were deployed on a cloud virtual machine (VM) on the Digital Research Alliance of Canada system. A Fed-BioMed client was deployed for each dataset involved in the federated learning experiments. The PPMI, ADNI, PREVENT-AD, QPN, Calgary, and mega-analysis datasets were deployed on the same machine at The Neuro’s Brain Imaging Centre. The COGTIPS dataset was deployed on a VM on the SURF Research Cloud system in the Netherlands.

## Funding acknowledgements

MW acknowledges support from the Canadian Institutes of Health Research (CIHR) (Canadian Graduate Scholarship – Doctoral 193412), the Brain Canada Foundation (Next Gen Award in Parkinson’s Disease Research), the Centre UNIQUE – Centre de recherche Neuro-IA du Québec (Doctoral Excellence Scholarship), the Fonds de Recherche du Québec – Santé (FRQS) (Bourse de doctorat en recherche BF2 329969), Parkinson Québec, and the Fonds de Recherche du Québec – Nature et technologie (FRQNT) (Bourse de maîtrise en recherche B1X 315843). The Brain Canada Next Gen Award in Parkinson’s Disease Research has been made possible by the Canada Brain Research Fund (CBRF), an innovative arrangement between the Government of Canada (through Health Canada) and Brain Canada Foundation, and by the Mireille and Steinberg Foundation and the Growling Beaver Brevet. The UNIQUE research centre is funded by the FRQNT Strategic Clusters Program.

JKP received funding from the Canadian Neuroanalytics Scholarship (CNS). The financial support of the Canadian Neuroanalytics Scholars Program, The Hilary & Galen Weston Foundation, and the support of Campus Alberta Neuroscience, the Hotchkiss Brain Institute at the University of Calgary, the Ontario Brain Institute and the Neuro at McGill University is greatly acknowledged.

GP-M received a Health-Professional Research Training Award from the Canadian Institutes of Health Research and a Research Fellowship from the Bay Tree Foundation and the Al Zaibak Family Fund.

JBP acknowledges support from the National Institutes of Health (NIH; NIH-NIBIB P41 EB019936 [ReproNim], NIH-NIMH R01 MH083320 [CANDIShare], NIH RF1 MH120021 [NIDM], and NIH-NIMH R01 MH096906 [Neurosynth]), the Canadian Institutes of Health Research (CIHR; PJT-185948, PJT-197805), the Michael J. Fox Foundation, the Quebec Parkinson Network, the McConnell Brain Imaging Centre, the Canada First Research Excellence Fund, awarded to McGill University for the Healthy Brains for Healthy Lives initiative (NeuroHub), the Chan Zuckerberg Initiative (EOSS5-0000000401), the Natural Sciences and Engineering Research Council of Canada (NSERC), The Galen and Hilary Weston Foundation (Canadian Neuroanalytic Scholar program and Big Bet Project), the Tanenbaum Open Science Institute, and the Brain Canada Foundation with support from Health Canada through the Canada Brain Research Fund in partnership with the Montreal Neurological Institute.

