## Supplementary information for "Open neuroinformatics infrastructure ecosystem for federated multisite studies"

| | N | Age (years) | Male | Female |
|---|---|---|---|---|
| **Age task** | | | | |
| *PPMI* | 1082 | 64.4 ± 8.1 | 524 (48.4%) | 558 (51.6%) |
| *Calgary* | 58 | 71.4 ± 7.3 | 28 (48.3%) | 30 (51.7%) |
| *QPN* | 69 | 62.6 ± 11.9 | 26 (37.7%) | 43 (62.3%) |
| *ADNI* | 315 | 74.4 ± 6.1 | 158 (50.2%) | 157 (49.8%) |
| *PREVENT-AD* | 342 | 64.8 ± 5.9 | 98 (28.7%) | 244 (71.3%) |
| *COGTIPS* | 29 | 62.3 ± 9.8 | 62.3 ± 9.8 | 12 (41.4%) |
| **Cognitive decline task** | | | | |
| *PPMI* | 768 | 62.8 ± 9.4 | 484 (63.0%) | 284 (37.0%) |
| *Calgary* | 61 | 70.6 ± 6.5 | 41 (67.2%) | 20 (32.8%) |
| *QPN* | 41 | 64.1 ± 8.5 | 29 (70.7%) | 12 (29.3%) |
| *ADNI* | 617 | 72.3 ± 7.4 | 325 (52.7%) | 292 (47.3%) |
| *PREVENT-AD* | 342 | 64.8 ± 5.9 | 98 (28.7%) | 244 (71.3%) |
| **Diagnosis task** | | | | |
| *PPMI* | 1908 | 63.7 ± 8.7 | 1044 (54.7%) | 864 (45.3%) |
| *Calgary* | 158 | 71.3 ± 6.7 | 94 (59.5%) | 64 (40.5%) |
| *QPN* | 290 | 64.8 ± 9.7 | 174 (60.0%) | 116 (40.0%) |
| *ADNI* | 1106 | 73.2 ± 7.1 | 586 (53.0%) | 520 (47.0%) |
| *COGTIPS* | 113 | 63.0 ± 8.1 | 68 (60.2%) | 45 (39.8%) |

**Supplementary Table 1: Participant demographics.** Age is given as mean ± standard deviation.

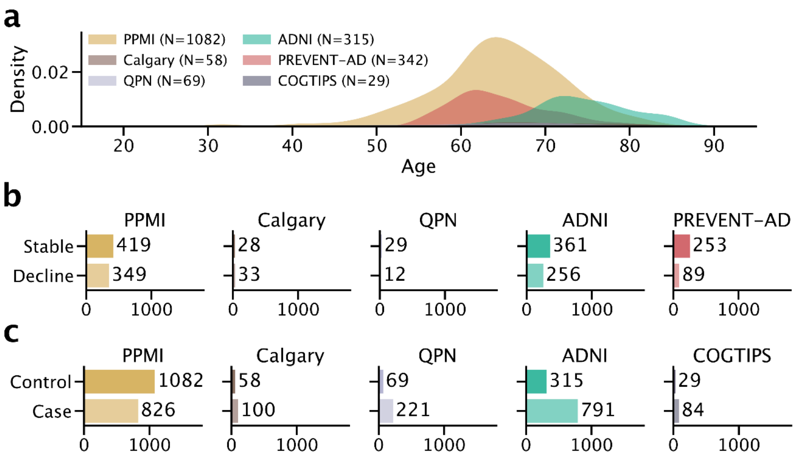


**Supplementary Figure 1: Distributions of target variables in prediction tasks. a**: Age. **b**: Cognitive decline. **c**: diagnosis.
